# O-GlcNAc regulates MTA1 transcriptional activity during breast cancer cells genotoxic adaptation

**DOI:** 10.1101/2021.02.08.430201

**Authors:** Xueqin Xie, Qiutong Wu, Keren Zhang, Yimin Liu, Nana Zhang, Qiushi Chen, Lingyan Wang, Wenli Li, Jianing Zhang, Yubo Liu

## Abstract

Chromatin modifier metastasis-associated protein 1 (MTA1), closely correlated with the development and progression in breast cancer, has a vital role in multiple cellular processes, including gene expression and cell homeostasis. Although MTA1 is a stress-responsive gene, its role in genotoxic adaptation remains unexplored. The current study sought to investigate the role of MTA1 and its O-GlcNAc modification in breast cancer cells genotoxic adaptation by using quantitative proteomics, ChIP-seq, transcriptome analysis, loss-and gain-of-functions experiments. We demonstrate that O-GlcNAc modification promotes MTA1 to interact with chromatin and regulates target gene expression, contributing to breast cancer cell genotoxic adaptation. MTA1 is modified with O-GlcNAc residues at serine 237/241/246 in adriamycin adaptive breast cancer cells and that modification improves the genome-wide interactions of MTA1 with gene promotor regions by enhancing its association with nucleosome remodeling and histone deacetylation (NuRD) complex. Further, O-GlcNAc-modulated MTA1 chromatin-binding influences the specific transcriptional regulation of genes involved in the adaptation of breast cancer cells to genotoxic stress. Our findings reveal a previously unrecognized role of O-GlcNAc MTA1 in transcriptional regulation and suggest that O-GlcNAc modification is a promising therapeutic target to overcome chemoresistance in breast cancers.

## Background

An essential feature of cancer cells is their ability to counteract various hostile environmental stresses, including nutrient deprivation, hypoxia and exposure to genotoxic agents [1]. Cancer cells develop mechanisms to reprogram stress response gene expression, evade death and assist cells escape from hypoxia and anti-cancer therapies [2]. Among the specialized pathways involved in cancer cells stress adaptation, chromatin modifier MTA1 (metastasis-associated protein 1) is recently proposed as a master regulator in stress response [3-5]. As a component of the nucleosome remodeling and histone deacetylation (NuRD) complex, MTA1 is one of the most upregulated oncogenes in human tumor [6]. It is widely reported that MTA1 modulates breast cancer metastasis and participate in gene transcriptional regulation through the chromatin remodeling of the target genes [7]. Although the *MTA1* was initially cloned as a metastasis-associated gene, its dual coregulatory activity of transcription was revealed until this regulator was detected in NuRD complex. Due to the interaction with diverse components of NuRD chromatin remodeling complex, MTA1 can act both as a coactivator or corepressor to regulate target gene transcription. Additionally, MTA1 is a stress-responsive gene and its accumulation in cancer cells has been linked with the resistance to multiple therapeutic agents, including genotoxicity drugs [6, 8]. Despite the intensive study on MTA1, the mechanism by which MTA1 modulates genotoxic adaptation and promotes tumor progression has not been clearly demonstrated.

Studies have shown that the regulation of MTA1 activity is achieved through at least two mechanisms: posttranslational modifications (PTM) of MTA1 [6] and interactions with its binding partners, including NuRD complex [9]. Post-translational addition of O-linked N-acetylglucosamine (O-GlcNAc) to protein serine and/or threonine is implicated as a key regulator of a wide range of nuclear and cytoplasmic proteins [10]. Two enzymes (O-GlcNAc transferase, OGT and O-GlcNAcase, OGA) dynamically catalyze this monosaccharide modification. As the major class of proteins regulated by O-GlcNAc modification, chromatin-binding proteins accounts for a quarter of all the O-GlcNAc modified proteins [11]. Aberrant O-GlcNAc modification of these proteins have been demonstrated to influence the transcriptional activity, DNA-binding and protein-protein interactions [12]. Therefore, this modification may affect multiple cancer-relevant genes transcription, and ultimately lead to tumorigenesis and progression. Recently, an association between OGT and the component NuRD complex, in particular MTA1, was identified by mass spectrum analysis [13]. This implies that OGT and O-GlcNAc modification may involve in modulating the transcriptional activity of MTA1 and NuRD complex. However, it is currently unknow whether MTA1 is O-GlcNAc modified and how the complex interplay between MTA1 and OGT affects the MTA1-modulated genotoxic adaptation in cancer cells.

In this study, we found that OGT interacts with MTA1 in breast cancer cells and promotes MTA1 O-GlcNAc modification at three serine (S) residues S237/S241/S246. We revealed that O-GlcNAc at these sites enhances the interaction between MTA1 and other NuRD components, in turn promotes the adaptation of breast cancer cells to genotoxic agent adriamycin (Adm). Further, genome-wide analysis of MTA1 chromatin-binding sites demonstrated that O-GlcNAc drives MTA1 to occupy gene promotor regions during genotoxic adaptation. We also provide evidence showing that O-GlcNAc-modulated chromatin occupation of MTA1 affect downstream genes expression and protect breast cancer cells against genotoxicity. Our results provided insights into the regulatory mechanisms of O-GlcNAc MTA1 and revealed that aberrant MTA1 O-GlcNAc modification plays an important role in cancer cells genotoxic stress adaptation.

## Results

### Elevated MTA1 expression level correlates with poor clinical prognosis in breast cancer

To investigate the potential role of MTA1 in breast cancer, we first analyzed *MTA1* in publicly available breast cancer and the normal tissue mRNA expression data from the Gene Expression Omnibus (GEO). We found that *MTA1* expression was significantly up-regulated in breast cancer samples (n = 30) compared with normal tissues (n = 9, *p* < 0.001, GSE1477, Figure 1a). Meanwhile, a strong association between high *MTA1* expression and shorter overall survival was observed when *MTA1* mRNA expression was compared in three GEO datasets (GSE16446, hazard ratio = 2.8, logrank *P* = 0.011; GSE9195, hazard ratio = 6.02, logrank *P* = 0.008; GSE31519, hazard ratio = 2.41, logrank *P* = 0.04), suggesting that accumulation of MTA1 may result in increased tumor aggressiveness (Figure 1b). Notably, we revealed a positive correlation between gene expression of *MTA1* and *OGT* with a moderate Pearson correlation coefficient (PCC) R value of 0.54 in breast cancer and breast tissue from the cancer genome atlas (TCGA) and the genotype-tissue expression (GTEx) datasets (Figure 1c). These results above indicated that MTA1 may be potentially regulated by OGT and play a role in breast cancer progression.

**Figure 1.**
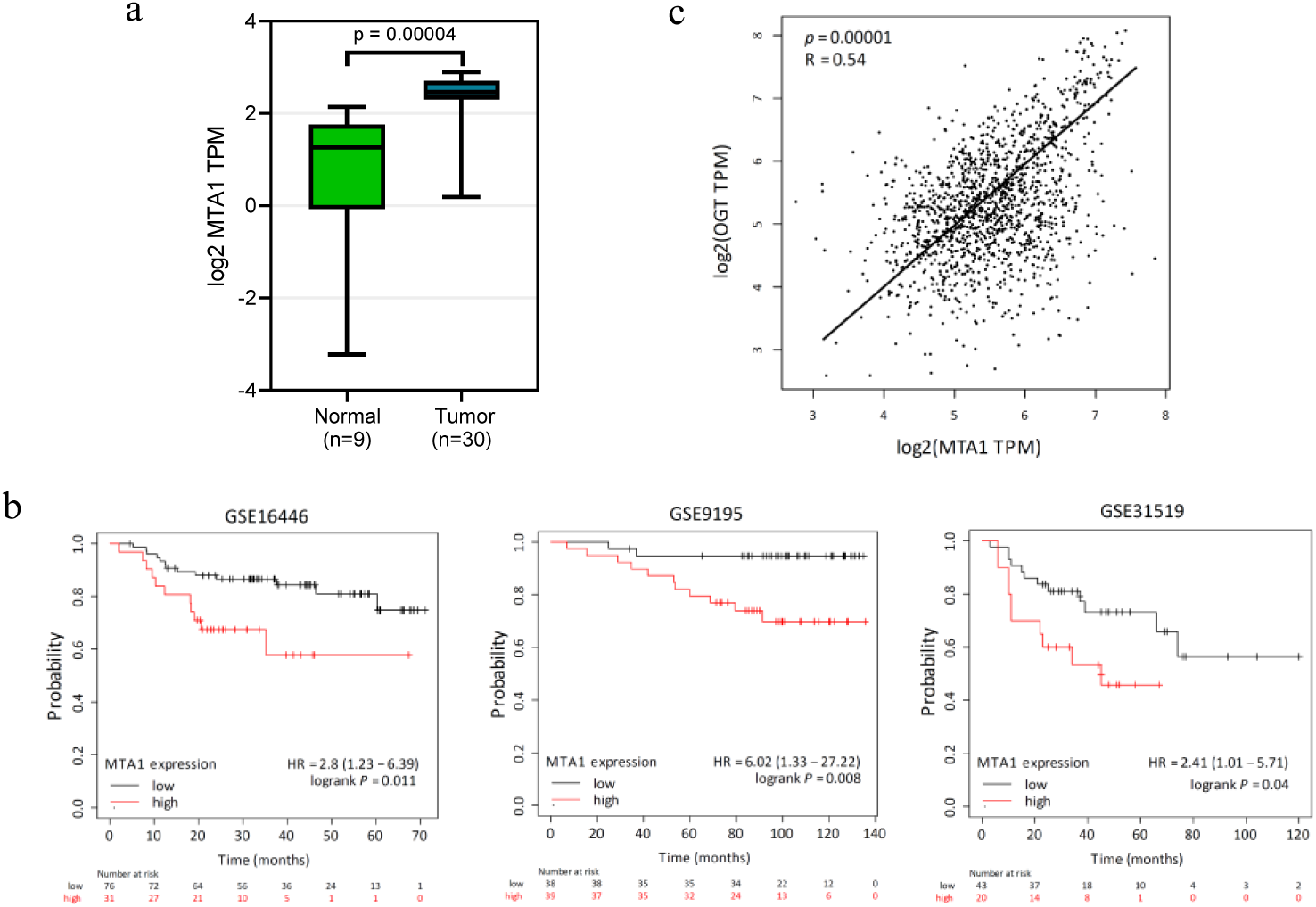
The expression of *MTA1* associates with poor clinical prognosis in breast cancer. (a) Transcriptional levels of MTA1 in breast cancer tissues and adjacent nontumoral breast tissues from the GEO database (GSE1477). The box plots show the medians (black lines), 25th and 75th percentiles (boundaries), and minimum/maximum values (whiskers). The *p* value is indicated. (b) *MTA1* positively correlates with overall survival in human breast cancer tissues. Datasets used for overall survival analysis are from indicated GEO datasets. Red lines represent the survival probability of patients with cancer with high *MTA1* expression levels and black lines represent that with low *MTA1* expression levels. HR, hazard ratio. The *p* values are indicated. (c) PCC R value revealed a moderate positive correlation between *MTA1* and OGT in TCGA and GTEx datasets samples.

### MTA1 binds to OGT and promotes the adaptation of breast cancer cells to genotoxicity

To further understand the mechanistic role of MTA1 and OGT in breast cancer, we explored their expression in a panel of breast cancer cell lines with different adaptability of genotoxic agent Adm. The immunoblotting results showed that cells with acquired (MCF-7/ADR) and intrinsic (MDA-MB-231) Adm adaptation contained obviously higher levels of MTA1 proteins than ADR sensitive cells (MCF-7 and T47D, Figure 2a, Supplementary Figure 1). Meanwhile, breast cancer cells also displayed a moderate increase in OGT protein levels upon the increasing of genotoxic adaptation, indicating the regulatory relationship between MTA1 and OGT. Analysis of the partners that interacted with MTA1 in NuRD complex, HDAC1, MBD3 and CHD4, revealed invisible correlation between the expression levels of these proteins and Adm adaptation. Further, MTA1 were also found to be O-GlcNAc modified in breast cancer cells. Co-immunoprecipitation (co-IP) of these cells showed that the protein-protein interactions (PPIs) between MTA1 and OGT were increased in Adm adaptive MDA-MB-231 and MCF-7/ADR cells compared with parental MCF-7 cells (Figure 2b). The interaction between MTA1 and other NuRD components revealed the same changing tendency, suggesting that O-GlcNAc might modulate the interaction between MTA1 and NuRD constituent proteins, in turn influences the genotoxic adaptation of breast cancer cells.

**Figure 2.**
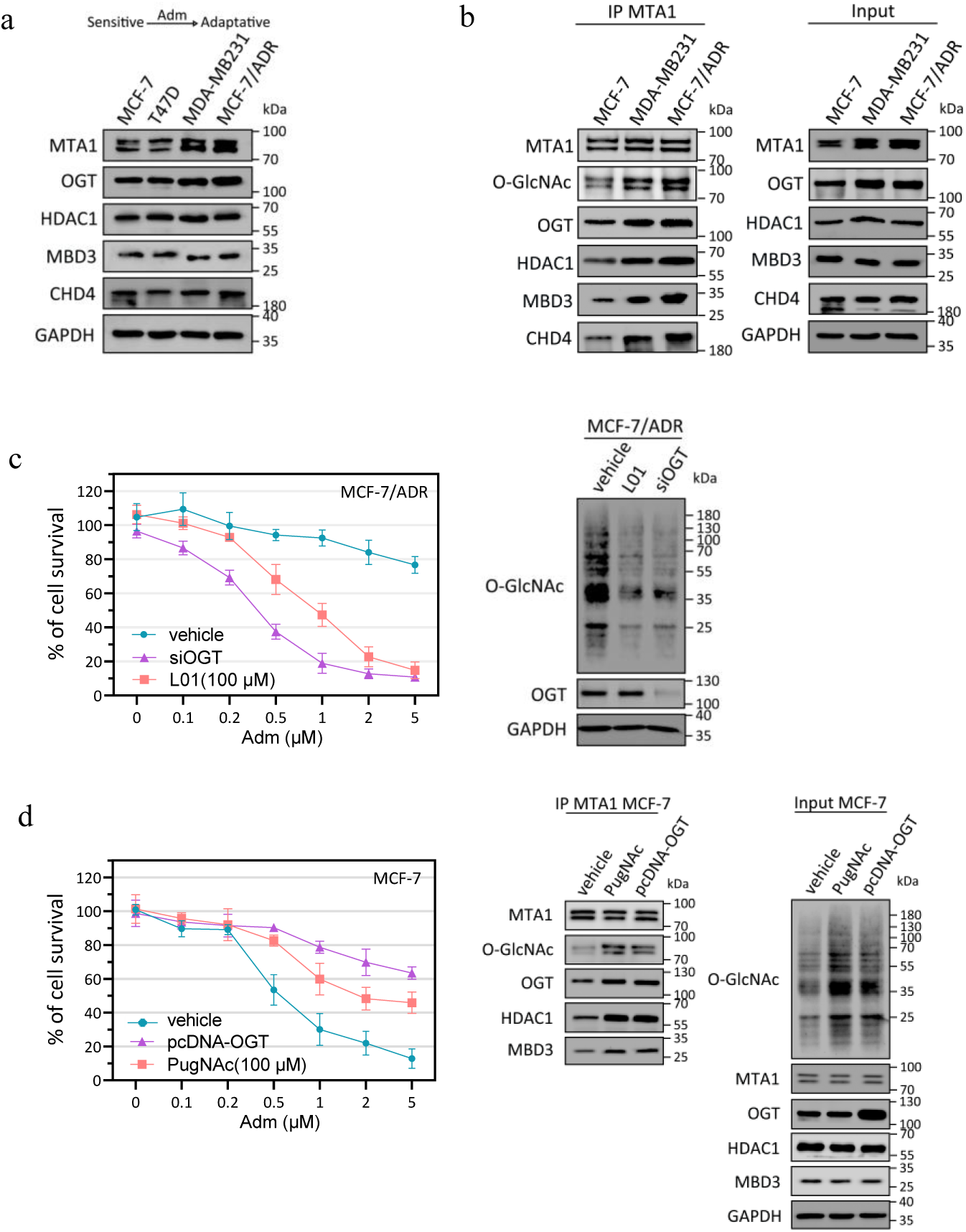
The interaction between MTA1 and OGT promote the adaptation of breast cancer cells to Adm. (a) Breast cancer cells with intrinsic and acquired Adm adaptation contain higher levels of MTA1 and OGT compared with Adm sensitive cells. The expression of indicated proteins in breast cancer cells were analysed by immunoblotting. (b) The PPIs between MTA1, OGT and NuRD components are increased in MDA-MB-231 and MCF-7/ADR cells compared with MCF-7 cells. MTA1 immunoprecipitation was performed and the immunoprecipitated fractions were analysed by immunoblotting by indicated antibodies. (c) The inhibition of O-GlcNAc modification increases the sensitivity of breast cancer cells to Adm and restrains the interaction of MTA1 with OGT and NuRD complex. MCF-7/ADR cells were transfected with OGT siRNA (siOGT) or treated with 100 μM L01 and then incubated with indicated doses of Adm for 48 h. The protein levels were examined by immunoblotting. Vehicle, cells treated with DMSO. The cell viability was assessed through an CCK8 assay. (d) The activation of O-GlcNAc modification increases the adaptation of MCF-7 cells to Adm and elevates the interaction of MTA1 with OGT and NuRD complex. MCF-7 cells were transfected with OGT expression plasmid (pcDNA-OGT) or treated with 100 μM PugNAc and then incubated with indicated doses of Adm for 48 h. The protein levels were examined by immunoblotting. Vehicle, cells treated with DMSO. The cell viability was assessed through an CCK8 assay.

Further, our results showed that the inhibition of O-GlcNAc modification with L01 restrained MTA1 from binding to OGT and NuRD complex. Similar results were obtained by silencing OGT with small interfering RNA (siRNA). O-GlcNAc inhibiting MCF-7/ADR cells showed increased sensitivity to Adm (Figure 2c). In contrast, OGA inhibitor PugNAc treatment or OGT overexpression in MCF-7 cells increased the interaction of MTA1, OGT and NuRD complex, subsequently elevated genotoxic adaptation (Figure 2d). In summary, these data indicated that O-GlcNAc modification could modulate the biological function of MTA1 and protects cancer cells from genotoxicity-induced death.

### MTA1 is O-GlcNAc modified at serine residues that enhances its interaction with NuRD complex

Next, we used a metabolic labeling (GalNAz) based O-GlcNAc sites identification and quantification proteomics method to compare the O-GlcNAc chromatin of MCF-7 and MCF-7/ADR cells (the workflow of this strategy is illustrated in Figure 3a). The O-GlcNAc chromatin-associated proteins can be metabolically labeled with azides, followed by reaction with a cleavable DADPS alkyne-biotin linker and enrichment with streptavidin beads. After cleavage of DADPS by formic acid, O-GlcNAc modified peptides were quantitative analysis using tandem mass spectroscopy (MS). Label free quantitative proteomics comparisons between biological replicates of MCF-7 and MCF-7/ADR O-GlcNAc chromatin revealed that 2033 O-GlcNAc chromatin-associated proteins exhibited ≥ 2-fold differences (*p* value ≤ 0.05, Figure 3b, Supplementary Table 1). Strikingly, OGT and chromatin-associated NuRD constituent proteins, including MTA1, were found to increase in genotoxic adaptative MCF-7/ADR cells compare with parental MCF-7 cells. Next, we explored MTA1 O-GlcNAc modified peptides in MS and the O-GlcNAc sites could be speculated. Combining the analysis by YinOYang (v1.2, online program for predicting potential O-GlcNAc sites), peptide QPSLHMSAAAASR spanning residues 235-247 was proposed as the O-GlcNAc modified MTA1 peptide (Figure 3c, Supplementary Figure 2).

**Figure 3.**
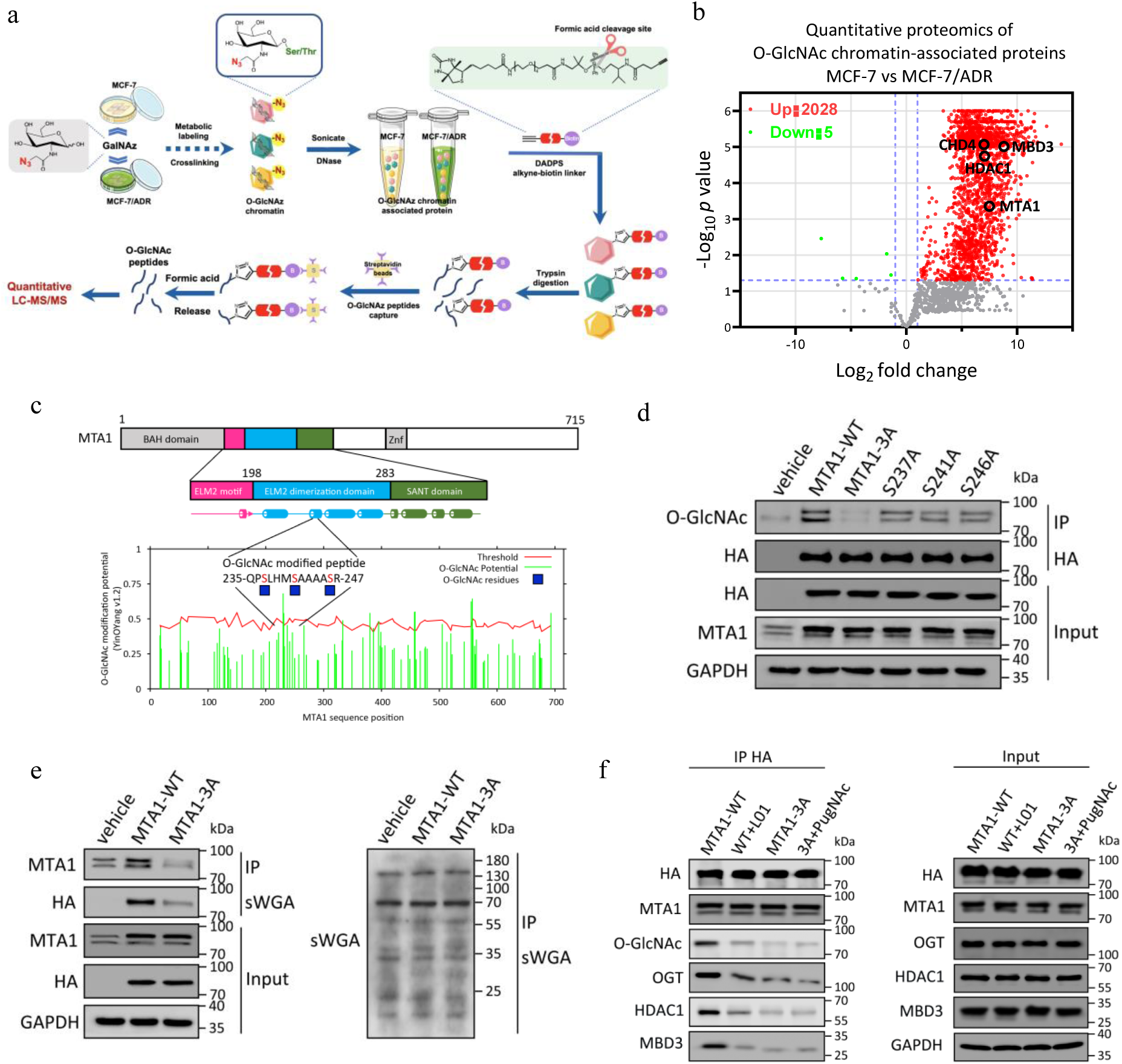
O-GlcNAc modification enhances the interaction of MTA1 with NuRD complex. (a) Schematic of the metabolic labeling (GalNAz) based O-GlcNAc sites identification and quantification proteomics method. MCF-7 and MCF-7/ADR cells were incubated on 1mM GalNAz media for 24 h. Chromatin was crosslinked and conjugated to a cleavable DADPS alkyne-biotin linker by a downstream click reaction. Biotin labeled chromatin proteins were enriched with streptavidin beads and subsequently digested with Trypsin. After cleavage of DADPS by formic acid, O-GlcNAc modified peptides were quantitative analysis using quantitative LC-MS/MS. (b) Volcano plot of quantitative proteomics data of O-GlcNAc chromatin-associated proteins in MCF-7 and MCF-7/ADR cells (n = 5 biologically independent experiments). The component of NuRD complex mentioned in this study are labeled. (c) Schematic representation of full-length MTA1 and its potential O-GlcNAc sites (predicted by YinOYang 1.2 server). (d) MTA1 is O-GlcNAcylated at S237/S241/S246. MCF-7/ADR cells expressing HA-WT-MTA1, HA-MTA1-S237A, HA-MTA1-S241A, HA-MTA1-S246A, or HA-MTA1-S237A/S241A/S246A (HA-MTA1-3A) were immunoprecipitated with anti-HA magnetic beads. O-GlcNAc modification was analyzed by immunoblotting. Vehicle, cells transfected with empty vector. (e) The O-GlcNAc levels of MTA1 is decreased after O-GlcNAc sites mutation. Immunoprecipitation was performed using O-GlcNAc-binding lectin sWGA, and the immunoprecipitated fractions were analyzed by immunoblotting for MTA1. Vehicle, cells transfected with empty vector. (f) After treatment with 100 μM L01 or PugNAc for 24 h, MCF-7/ADR cells expressing HA-MTA1-WT or HA-MTA1-3A were immunoprecipitated with anti-HA magnetic beads. Indicated proteins were analyzed by immunoblotting.

To further determine the potential O-GlcNAc sites, we generated a series of mutated MTA1 construct, in which the three serine residues within the above peptide were mutated to alanine (A) (HA-MTA1-S237A, HA-MTA1-S241A, HA-MTA1-S246A, HA-MTA1-S237A/S241A/S246A(HA-MTA1-3A)). HA-tagged wild-type MTA1 (HA-MTA1-WT) and mutated MTA1 constructs were then stably transfected into MCF-7/ADR cells and their O-GlcNAc modification levels were compared. As shown in Figure 3d, the levels of MTA1-3A O-GlcNAc modification (O-GlcNAc specific antibody CTD110.6) was dramatically decreased compared to that of HA-MTA1-WT, while single site mutants showed a slightly diminished level of O-GlcNAc MTA1, supporting the notion that the majority of MTA1 O-GlcNAc modification occur on S237/S241/S246 in this genotoxic adaptative cell model. Similar results were observed using the O-GlcNAc-binding lectin succinylated wheat germ agglutinin (sWGA, Figure 3e). Our experiments also showed that low O-GlcNAc levels decreased the interaction of MTA1 with OGT and NuRD complex in MCF-7/ADR cells (Figure 3f). Therefore, O-GlcNAc modification on serine residues could enhance the function of MTA1 through modulating the constituent of NuRD complex.

### O-GlcNAc improves the genome-wide interactions of MTA1 with gene promotor regions in the response to genotoxic stress

Since O-GlcNAc MTA1 have been implicated in genotoxicity adaptation and the emerging roles of MTA1 have been highlighted in gene transcriptional regulation, we wonder whether O-GlcNAc modulates the DNA binding feature of MTA1. To this end, we performed chromatin immunoprecipitation followed by sequencing (ChIP-seq) using anti-HA magnetic beads in HA-MTA1-WT or HA-MTA1-3A stably expressed MCF-7 cells (GSE162932). The ChIP-seq peaks of MTA1-WT and MTA1-3A showed distinguishing characteristics across the genome (Figure 4a). Among these binding sites of MTA1-WT, 17.6% were located in the promotor regions, while only 4.1% MTA1-3A peaks distributed to the transcription start sites (TSS), indicating that O-GlcNAc modification induces a genome-wide promoter-biased association of MTA1 in breast cancer cells (Figure 4b). Consisting with this, the ChIP-seq signals at TSSs obtained for MTA1-WT was remarkably stronger than that found for MTA1-3A (Figure 4c). To confirm O-GlcNAc-induced ChIP-seq distribution to promoters, we measured the histone markers at these MTA1 binding sites using published ChIP-seq datasets generated from MCF-7 cells. The MTA1-WT binding sites shared a high degree of conservation with transcriptional activation mark histone H3 acetylated at lysine 27 (H3K27ac), whereas a low signal for histone H3 trimethylated at lysine 27 (H3K27me3, repressive mark) was found [14]. However, low H3K27ac and high H3K27me3 signal were observed in MTA1-3A peaks. MTA1-WT located regions were surrounded by higher levels of histone H3 trimethylated at lysine 4 (H3K4me3, promoter mark) compared with that of MTA1-3A (Figure 4d). These data indicate that O-GlcNAc-modulated chromatin occupation plays a predominant regulatory role in MTA1 transcriptional activity regulation.

**Figure 4.**
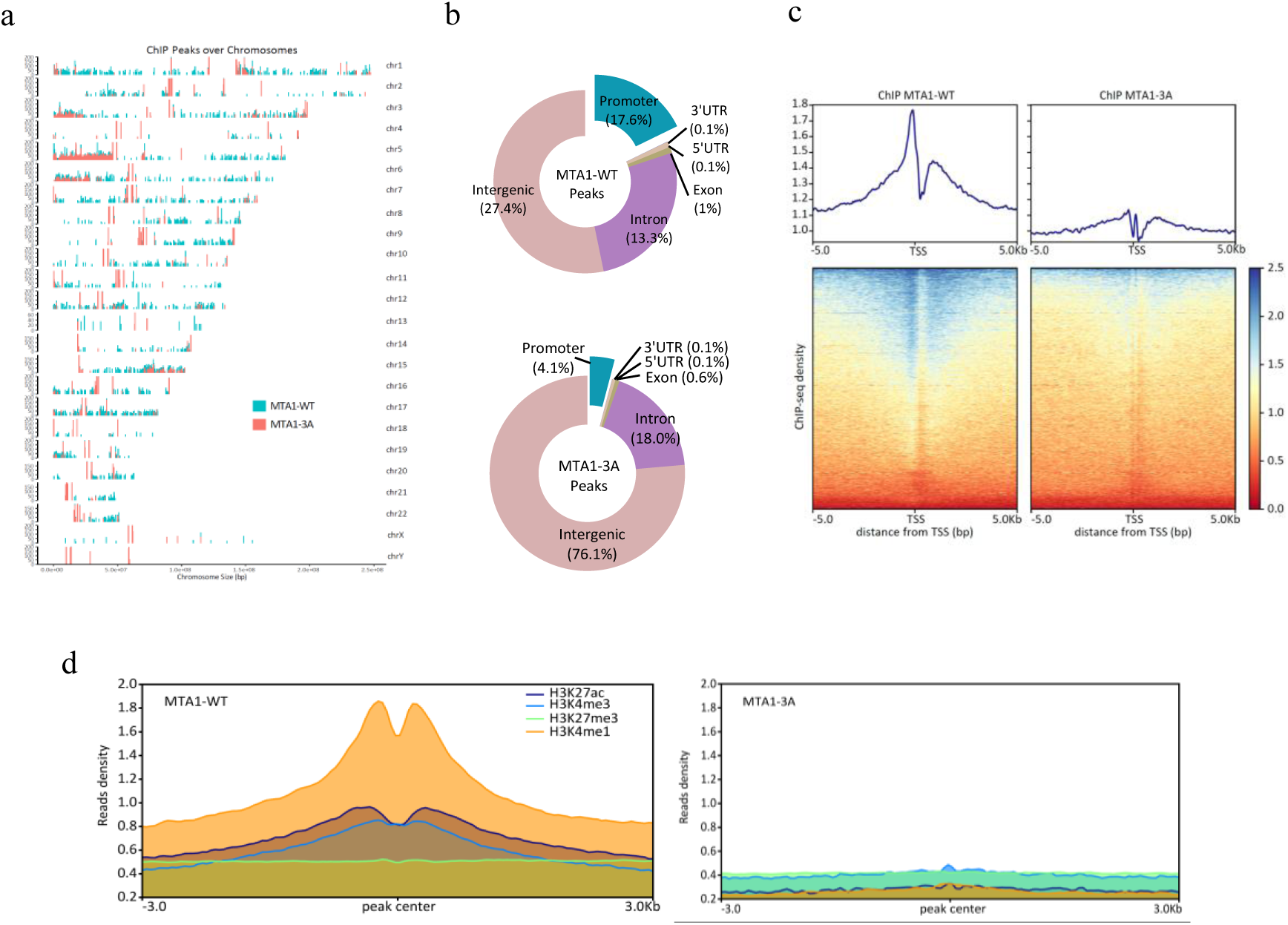
O-GlcNAc influences the genome-wide distributions of MTA1 on the chromatin. (a) Coverage plot of the MTA1 ChIP-seq peak locations over the whole human genome. The peaks of HA-MTA1-WT and HA-MTA1-3A stably expressed MCF-7 cells showed distinct characteristic. (b) HA-MTA1-WT and HA-MTA1-3A ChIP-seq peak annotation relative to known genomic elements. (c) Heat maps of the ChIP-seq signal density at TSSs (±3 kb). The average signal profile is shown. The red color indicates low a signal, and a blue color indicates a high signal. (d) Average enrichment profiles of published H3K27ac, H3K4me3 (GSE97481), H3K27me3 (GSE96363) and H3K4me1 (GSE86714) ChIP-seq reads at HA-MTA1-WT and HA-MTA1-3A ChIP-seq peaks.

### O-GlcNAc-modulated chromatin occupation of MTA1 contributes to stress adaptive genes transcription

To further illustrate the underlying mechanism by which O-GlcNAc MTA1 regulate breast cancer cells genotoxic adaptation, we performed quantitative comparisons of the peaks from MTA1-WT and MTA1-3A ChIP-seq datasets using MAnorm software [15]. A great part of peaks (unbiased peaks) was concurrent between the above datasets, while 4574 differential quantitative peaks were identified (MTA1-WT-biased peaks, Figure 5a). Correspondingly, the differential quantitative peak associated genes were categorized based on Gene Ontology (GO) terms related to the cell adhesion and mobility, cellular response to drug, transcription factor activity, tumor necrosis factor (TNF) signaling pathway, and apoptosis (Figure 5b). Subsequent gene expression analysis by RNA-seq identified 950 differentially expressed genes (DEGs, false discovery rate (FDR) ≤ 0.05 and fold change ≥ 1.5) between MTA1-WT and MTA1-3A stably expressed MCF-7 cells (Figure 5c, Supplementary Table 2, GSE162932). GO and gene set enrichment analysis (GSEA) revealed that DEGs were enriched in TNF production and secretion pathway (Figure 5d), indicating that O-GlcNAc-modulated fluctuations of MTA1 chromatin occupation regulated these genes transcription and adjust the cells to genotoxic insult. For the integrated assessment of O-GlcNAc MTA1-regulated gene expression, the differentially quantitative MTA1 ChIP-seq peak-associated genes (Supplementary Table 3) were matched to the above RNA-seq DEGs and yielded 95 overlapping genes (Figure 5e, Supplementary Table 4). Of note, the majority of genes were enriched in the cellular stress response, TNF signaling pathway and cell adhesion. We further confirmed that MTA1 knockdown MCF-7/ADR cells were more sensitive to Adm treatment compared with the viability of control cells also treated with Adm (Figure 5f). MTA1-3A-transfected MCF-7/ADR cells were more sensitive to Adm than MTA1-WT-transfected cells, suggesting the protective role of O-GlcNAc MTA1. Together, O-GlcNAc-modulated MTA1 chromatin binding participates in the expression of stress adaptive genes.

**Figure 5.**
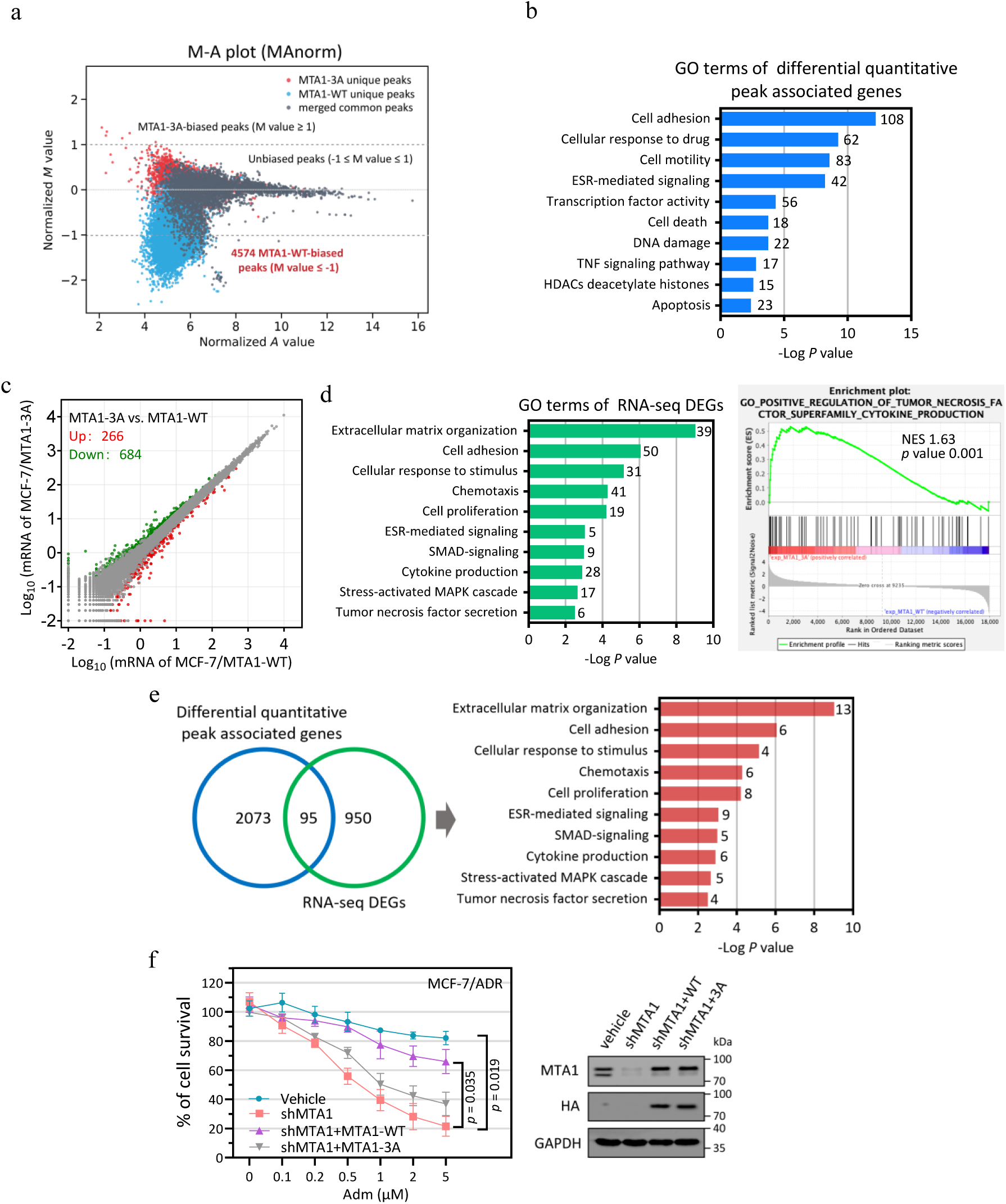
O-GlcNAc MTA1 contributes to stress adaptive genes expression. (a) Normalization and quantitative comparisons of ChIP-seq peaks using MAnorm. MA plots of all peaks from both HA-MTA1-WT and HA-MTA1-3A ChIP-seq datasets after MAnorm. Differential quantitative peaks (“MTA1-WT-biased” or “MTA1-3A-biased”, M value (log2 fold change) ≥ 1 or ≤ -1 and *P* value ≤ 10^−5^) are shown. (b) The differential quantitative peak associated genes were used for GO term analysis. (c) Scatter plot showing the RNA-seq DEGs expression levels (fragments per kilobase of transcript per million mapped reads (FPKM), fold change ≥ 1.5, FDR ≤ 0.05) in HA-MTA1-WT expressed MCF-7 cells compared with HA-MTA1-3A expressed MCF-7 cells. n = 2 biologically independent RNA-seq replicates. (d) Left images: GO analysis of RNA-seq DEGs in HA-MTA1-WT and HA-MTA1-3A expressed MCF-7 cells. The gene numbers identified in certain terms are shown. Right images: GSEA of RNA-seq DEGs. The normalized enrichment score (NES) and *p* value are indicated. (e) Overlap of 2073 differential quantitative peak associated genes identified from ChIP-seq with 950 DEGs in RNA-seq is shown in a Venn diagram. The 95 overlapping genes were used for GO term analysis. (f) The rescue expression of HA-MTA1-WT and HA-MTA1-3A were performed in MCF-7/ADR-shMTA1 cells (shMTA1+MTA1-WT and shMTA1+MTA1-3A, respectively). Cells were subsequently treated with increasing doses of Adm for 48 h. Vehicle, cells transfected with empty vector. The cell viability was assessed using a CCK8 assay. The protein levels of MTA1 were analyzed by immunoblotting.

### O-GlcNAc MTA1 play a key role in the transcriptional activation of c-Fos to protect breast cancer cells against genotoxicity

To corroborate the role of O-GlcNAc in modulating MTA1 transcriptional activity, we focused on a novel O-GlcNAc MTA1 promoted gene, *c-Fos*, which was reported to participate in regulating TNF signaling [16]. As shown in Figure 6a, the promoter regions of *c-Fos* were robustly bound by MTA1-WT but weakly bound by MTA1-3A in MCF-7 ChIP-seq datasets. ChIP-qPCR assays also confirmed the interaction of O-GlcNAc MTA1 and the *c-Fos* promoter in MCF-7 cells. O-GlcNAc sites mutation significantly reduced the binding of MTA1 with *c-Fos* promoter (two-sided unpaired Student’s t-test, *p* = 0.006). We next utilized luciferase reporter system to test the affection of O-GlcNAc on the transcriptional efficiency of MTA1 in MCF-7/ADR cells. As shown in Figure 5b, O-GlcNAc elimination by sites mutation and L01 treatment significantly decreased the expression of the reporter harboring the promoter region of *c-Fos*. Consistently, we confirmed that MTA1-mediated increase in the mRNA and protein levels of c-Fos was dependent on O-GlcNAc modification (Figure 6c). MTA1 knockdown by shRNA significantly inhibited c-Fos expression in both mRNA level in MCF-7/ADR cells. The expressions of c-Fos could be restored by MTA1-WT rescue. Notably, MTA1-3A rescue or L01 treatment was significantly less effective to increase the transcription of *c-Fos*. O-GlcNAc inhibition of MTA1 also suppressed the protection against Adm stress in MCF-7/ADR cells. Furthermore, c-Fos knockdown using siRNA obviously reduced the protection of MCF-7/ADR cells from Adm, which indicates the adaptive role of this gene in genotoxicity (Figure 6d). These results provide direct evidence that activation of MTA1 by O-GlcNAc modification modulates downstream gene expression to protect breast cancer cells against genotoxicity.

**Figure 6.**
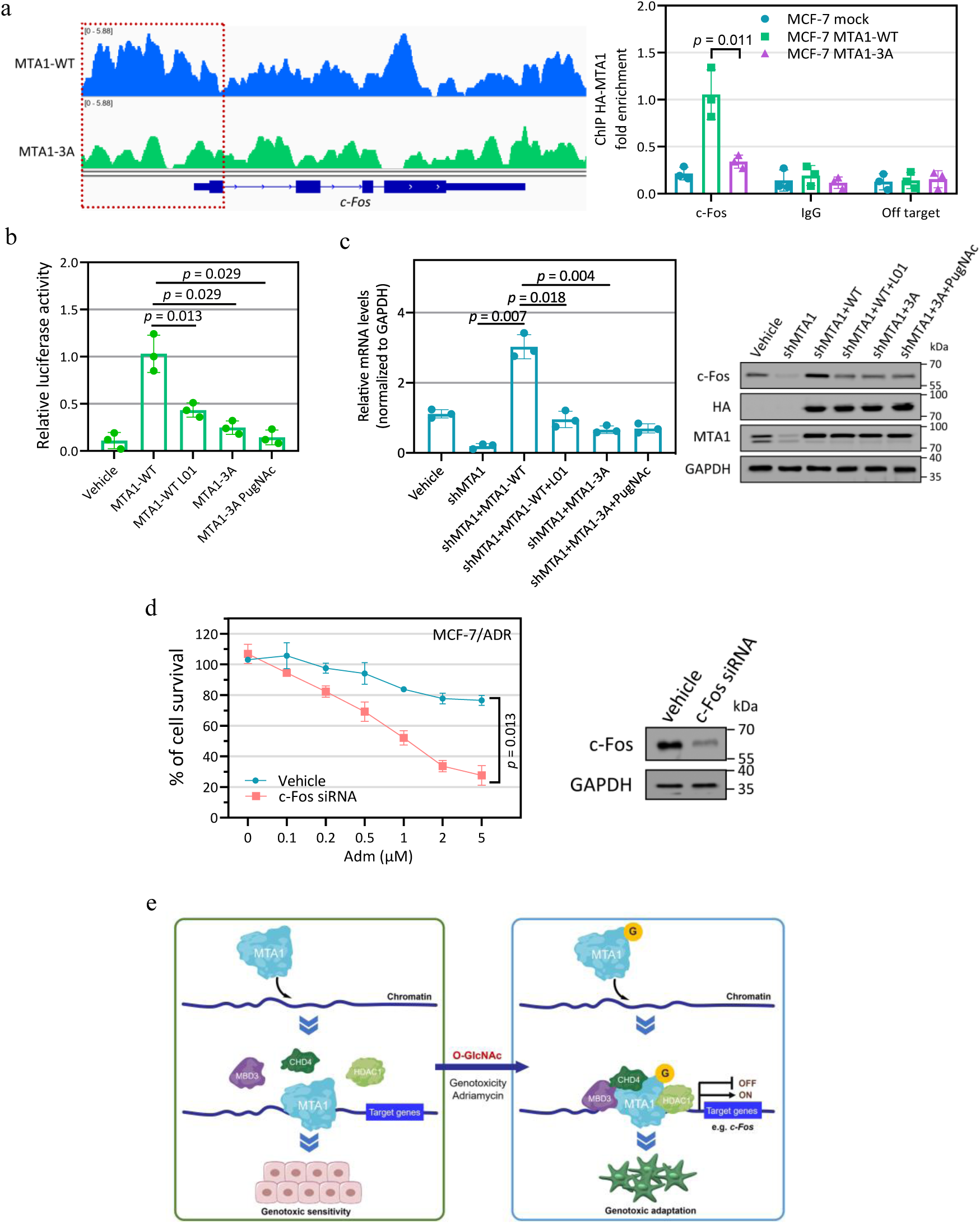
O-GlcNAc MTA1 activates the transcription of *c-Fos* and protects breast cancer cells against genotoxicity. (a) Left images: Integrative Genomics Viewer (IGV) tracks showing ChIP-seq signal at the promoter regions of *c-Fos*. Right images: Validation of O-GlcNAc MTA1 binding peaks by ChIP-qPCR. Each bar represents the fold enrichment of binding relative to the input. IgG and random primers that could not specifically bind the indicated gene promoter regions (off target) were used as negative controls. Mock, MCF-7 cells transfected with empty vector. (b) MCF-7/ADR cells were transfected with a reporter vector consisting of luciferase cDNA fused to the *c-Fos* promoter. The pGL3-basic vector (Vehicle) was used as a control. (c) Effect of MTA1 O-GlcNAc modification on *c-Fos* transcription and protein levels in MCF-7/ADR cells. The gene mRNA levels were analyzed by qPCR. The protein levels were analyzed by immunoblotting. Vehicle, cells transfected with empty vector. (d) MCF-7/ADR cells were transfected with c-Fos siRNA and then incubated with indicated doses of Adm for 48 h. Vehicle, cells treated with DMSO. The cell viability was assessed through an CCK8 assay. The protein levels were examined by immunoblotting. (e) Proposed model for the MTA1 transcriptional activity regulated by O-GlcNAc modification during breast cancer cells genotoxic adaptation. O-GlcNAc modification enhances the interaction of MTA1 with NuRD complex and chromatin. O-GlcNAc MTA1 preferentially modulates target genes transcription and drives breast cancer cells adaptation during genotoxic stimulation.

## Discussion

Breast cancer is the most common malignancy in the female population and has presently afflicted more than 2.5 million women worldwide [17]. Although therapies for breast cancer have been greatly improved, resistance to chemotherapy, including genotoxic agents, remains a paramount challenge in the treatment of this tumor [18]. However, the exact mechanisms behind the drug resistance are still poorly understood. One of the possible mechanisms of drug tolerant origin is the adaptive phenotypic plasticity of cancer cells [19]. Genotoxic drug exposure results in dynamical cellular stress response that modulates the gene expression involves in DNA damage repair or cell death inhibition, which follows by re-establishment of cellular homeostasis and gives rise to cancer cells genotoxicity adaptation [20]. This dynamic transcriptional regulation requires the recruitment and coordinated action of chromatin modifiers, such as MTA1-containing NuRD complex [21].

MTA1 is a cancer progression-related gene product that was initially isolated from invasive breast cancer cells [7]. It was later proved that MTA1 is a stress response protein that its expression can be acutely upregulated upon the treatment of various stress agents [3]. A growing body of evidence has shown that MTA1 modulates the transcription of estrogen receptor and promotes the progression and drug resistance of breast cancer [22-24]. MTA1 and its interacting proteins have also been found to regulate the transcription of estrogen-regulated genes and contribute to the development of hormone independence of breast cancer cells. This suggests that MTA1-modulated transcriptional regulation of gene expression programs could activate the cell survival pathways for counteracting emerging apoptotic pathways induced by stress stimulation. Findings presented here demonstrated that MTA1 was significantly overexpressed in breast cancer patient samples compared with the normal breast tissues. We also found that high MTA1 mRNA levels were associated with adverse outcome in breast cancer patients, indicating MTA1 may have an important role in breast cancer progression. Since Adm is one of the most active chemotherapy agents in breast cancer treatment [25], we used this genotoxic agent to investigate the stress adaptive function of MTA1 in breast cancer cells. Cells with genotoxic adaptation were characterized by higher levels of MTA1 than sensitive breast cancer cells. Notably, the interaction between MTA1 and OGT were increased in Adm resistance cells, suggesting that OGT and O-GlcNAc modification may have an inherent contribution of MTA1-dependent cell survival mechanisms of genotoxic adaptation in breast cancer cells.

The transcriptional activity of the MTA1 protein is largely regulated by its PTM [6]. For example, SUMOylation and methylation is necessary for the corepressor activity of MTA1, whereas acetylation plays a key role in modulating its transforming activity [26]. Here, our results revealed that O-GlcNAc modification of MTA1 enhances its ability to interact with NuRD components and contribute to the formation of functional transcriptional regulation complexes. Emerging evidence has indicated that O-GlcNAc modification has been recognized as a common feature of cancer cells and dynamically regulates the activities and functions of nuclear proteins to cope cellular stress [27]. Our previous studies have demonstrated that genotoxicity agents dynamically induce global O-GlcNAc elevation and widely influence the gene expression pattern in cancer cells [28]. Using the metabolic labeling-based O-GlcNAc sites identification and quantification proteomics method, we revealed that O-GlcNAc chromatin-associated proteins increase their interaction with the genomic DNA in genotoxic adapted MCF-7/ADR cells compare with parental MCF-7 cells. Further, the chromatin binding activity of MTA1 can be enhanced by O-GlcNAc modification. This could be due to the O-GlcNAc-induced recruitment of NuRD complex to MTA1. The majority O-GlcNAc sites of MTA1 is S237/S241/S246, which located in the egl-27 and MTA1 homology domain 2 (ELM2) domains. The crystal structure (PDB: 4BKX) of the complex between ELM2 domain of MTA1 and HDAC1 shows that the ELM2 domain positions close to HDAC1 and contributes to the homodimerization of HDAC1:MTA1 complexes. Considering that O-GlcNAc sites mutations separate MTA1 from NuRD components including HDAC1, it is therefore tempting to speculate that O-GlcNAc modification at these sites is likely required for the dynamic association/dissociation of NuRD complex and thereby transcriptional regulation of downstream genes. Although the majority of NuRD components were reported to interact with OGT recently [13], we identified them in metabolic labeling-based proteomics comparison with different quantitation. It will be of great value to examine whether O-GlcNAc modification of other NuRD components also affects their biological activity in future studies.

We next tested further whether the effect of O-GlcNAc sites mutations represent impaired DNA-binding functions of MTA1. Quantitative comparison between MTA1-WT and MTA1-3A ChIP-seq signals provide insight into the differences in O-GlcNAc-modulated chromatin elements occupancy of MTA1. We reveal a promoter aggregation in O-GlcNAc MTA1 chromatin-binding sites, suggesting that O-GlcNAc modification prompts MTA1 to reprogram the expression of genes needed for genotoxic adaptation. Combining the variation of O-GlcNAc-modulated MTA1 chromatin-binding sites with DEGs from transcriptome datasets of MTA1-WT and MTA1-3A expressed cells, a part of O-GlcNAc MTA1 target genes are upregulated, whereas the expression of others is attenuated, which agrees with the dual coregulatory activity of this chromatin modifier and reveals the complexity of O-GlcNAc-modulated gene transcription. A large fraction of O-GlcNAc MTA1 targets, including *c-Fos* are predicted to participate genotoxic stress response. Beyond these genes, several genes which were previously not reported to regulate cellular genotoxic resistance were further predicted in this study. Such future studies may include a functional verification of these newly discovered O-GlcNAc MTA1 target genes in breast cancer genotoxic adaptation.

In summary, our results demonstrate that O-GlcNAc modification at S237/S241/S246 in MTA1 enhances its interaction with NuRD complex and chromatin, thereby preferentially modulates target genes transcription to drive subsequent adaptation during genotoxic stimulation (Figure 6e). Base on the key role of MTA1 in breast cancer progression and genotoxic resistance, O-GlcNAc modification adds a further layer of regulation to this chromatin modifier. These evidences, combined with the observation that many chromatin-associated proteins including remodeling factors are O-GlcNAc modified, suggest the potential of O-GlcNAc modification as a crucial modulator of chromatin activity and function.

## Materials and methods

### Cell culture, reagents and transfection

MCF-7, T47D and MDA-MB-231 cells were obtained from Type Culture Collection of the Chinese Academy of Sciences (Shanghai, China) and were used within 6 months from resuscitation. All the cells were cultured in 90% RPMI-1640 (Gibco) supplemented with 1 % penicillin/streptomycin antibiotics (Gibco) and 10 % fetal bovine serum (Gibco). Adm (Adriamycin, Sigma) was added to the cell cultures in stepwise increasing concentrations from 0.1 to 10 μM for about eight months to develop genotoxicity adapted MCF-7/ADR cells. MCF-7/ADR cells were maintained in complete medium without Adm for one week and cells with > 90% viability before subsequent treatments. L01 was purchased from BioBioPha Co., Ltd. PugNAc were purchased from Sigma. Other reagents were used as analytic grade or better. Full-length wild type (WT) human MTA1 and O-GlcNAc amino acid sites mutant ((S237, S241, S246, S237/S241/S246)→A) MTA1 were subcloned into pCMV-Puro64. The primers are: sense-5’ ACCGAATTCGCCACCATGGCCGCCAACATGTACAGG 3’; antisense-5’ CTAGGATCCGTCCTCGATGACGATGGGC 3’. *c-Fos* promoter report constructs were made by cloning 2102 bp of human *c-Fos* promoter into luciferase vector pGL3-basic. The primers are: sense-5’ ATGTGTGTTTTT CCGTTTCCCTCCCT 3’; antisense-5’ GCTGTGGAGCAGAGCTGGGTAGGAG 3’. OGT siRNA (#sc-40781), c-Fos siRNA (#sc-29221) and MTA1 shRNA (#sc-35981-SH) were purchased from Santa Cruz Biotechnology. The construction of pcDNA-OGT was reported previously [28]. Transfection of MCF-7 and MCF-7/ADR cells was performed with Lipofectamine 3000 (Invitrogen) according to the manufacturer’s instructions. The stably transfected cells were then selected by the addition of puromycin (Sigma) to the medium.

### Immunoblotting/lectin blotting and co-immunoprecipitation

For immunoblots/lectinblots, the proteins were resolved on SDS-polyacrylamide gel electrophoresis gels followed by standard immunoblots. The primary antibodies used were anti-O-GlcNAc CTD110.6 (BioLegend, #838004), anti-OGT (Proteintech, #11576-2-AP), anti-GAPDH (CST, #5174), anti-HA (CST, #3724), anti-MTA1 (CST, #5646), anti-CHD4 (CST, #12011), anti-MBD3 (CST, #99169) and anti-HDAC1 (CST, # 34589). Lectin sWGA (Vector Laboratories, #B-1025S) was used for lectin blotting. The appropriate secondary antibody used were anti-mouse IgG-HRP (CST, # 7076), anti-rabbit IgG-HRP (CST, #7074), anti-mouse IgM-HRP (Abcam, #ab97230), and the signals were detected by the ECL Plus kit (GE Healthcare). All blots are representative of at least two independent experiments.

For co-immunoprecipitation, cells were harvested and lysed with western/IP lysis buffer (Beyotime, #P0013). Immunoprecipitates were washed five times and then subjected to immunoblotting analysis. The antibodies and beads used for IP were anti-MTA1 (CST, #5646), anti-HA-magnetic beads (Bimake, #B26201) and protein A/G-magnetic beads (Bimake, #B23201).

### Cell viability assay

Cells were seeded in a 96-well plate and treated with vehicle or the indicated doses of Adm with/without L01 or PugNAc for 48 h. The viable cells were determined by Enhanced Cell Counting Kit-8 (Beyotime, #C0041) according to the manufacturer’s instructions. 450 nm absorbance of each well was measured and normalized to the vehicle-treated wells.

### Metabolic labeling

The cells at 30% confluence in 10 cm dishes were incubated with culture medium containing 1 mM GalNAz for 48 h. 1 × 10^7^ cells were harvested by trypsin digestion and washed for three times with PBS and then crosslinked with 1% formaldehyde for 15 min. Crosslinking was quenched with 2.625 M glycine. The cells were scraped from plate and spun down and resuspended in 1 ml hypotonic buffer (10 mM HEPES (pH 7.5), 10 mM KCl, 0.1 mM MgCl_2_, 0.4% Igepal CA-630, Protease inhibitor cocktail (Roche) and phosphatase inhibitor cocktail (Roche)) until cell cytomembrane were broken. The nuclei was spun down and washed in hypotonic buffer four times, homogenized in 500 μL lysis buffer (50 mM HEPES (pH 7.5), 150 mM NaCl, 1.5 mM MgCl_2_, 1% Igepal CA-630, 0.1% SDS, protease inhibitor cocktail and phosphatase inhibitor cocktail) and incubated on a rotating stand for 1 h at 4°C. Lysed nuclei were treated with Benzonse Nuclease (Millipore, #70664) at 4°C for 1 h and centrifuged at 20,000 g for 15 min at 4°C. Supernatant was collected and incubated with 120 μM dialkoxydiphenylsilane (DADPS) alkyne-biotin (Click Chemistry Tools, #1331-25), 100 μM BTTAA (Click Chemistry Tools, #1236-500), 50 μM CuSO_4_ and 2.5mM freshly prepared sodium ascorbate at 25°C for 2 h. 10 mL methanol was added to the above solution and stored at -80 °C for 4 h. The precipitants were centrifuged at 10,000 g for 15 min at 4°C, washed twice with ic e-cold methanol, and dissolved in 1mL western/IP lysis. The resulting solution was pre-cleared with pre-cleared with 100 μL vehicle-magnetic beads (BEAVER Life Science), then 200 μL streptavidin-magnetic beads (BEAVER Life Science, #22308-10) were transferred into the above solution. The resulting mixture was incubated at 4°C for 4 h on a rotating stand. The beads were then washed with PBS for five times and Milli-Q water for five times.

The washed beads were resuspended in 500 µl PBS containing 6 M urea and 10 mM DTT and incubated at 35 °C for 30 min, followed by addition of 20 mM iodoacetamide for 30 min at 35 °C in the dark. The beads were the n collected by centrifugation and resuspended in 200 µL PBS containing 2 M urea, 1 mM CaCl _2_ and 10 ng/μL trypsin (Promega). Trypsin digestion was performed at 37 °C with rotation overnight and the beads were washed three times with PBS and distilled water again. The modified peptides were released from the beads with two treatments of 2% (v/v) formic acid/water (200 µL) for 1 h with gentle rotation, and the eluent was collected and combined. The beads were then washed with 50% (v/v) acetonitrile/water containing 1% formic acid (400 µL), and the washes were combined with the eluent to form the cleavage fraction. The labeled peptides were dried in a vacuum centrifuge and analysed by LC-MS/MS.

### LC-MS/MS analysis

Peptides were re-dissolved in solvent A (0.1% FA in 2% ACN) and directly loaded onto a trap column (Acclaim PepMap 100, Thermo Fisher Scientific). Peptide separation was performed using a reversed-phase analytical column (5 μm C18, 75 μm × 25 cm, home-made) with a linear gradient of 5–26% solvent B (0.1% FA in 98% ACN) for 85 min, 26-35% solvent B for 10 min and 35-80% solvent B for 8 min at a constant flow rate of 1 μL/min on an UltiMate 3000 RSLCnano system. The resulting peptides were analyzed by Q Exactive HF-X mass spectrometer (Thermo Fisher Scientific). The peptides were subjected to ESI source followed by tandem mass spectrometry (MS/MS) coupled online to the UHPLC (Thermo Fisher Scientific). Intact peptides were detected in the Orbitrap at a resolution of 60,000. Peptides were selected for MS/MS using 28% NCE; ion fragments were detected in the Orbitrap at a resolution of 15,000. A data-dependent acquisition procedure that alternated between one MS scan followed by 30 MS/MS scans was applied for the top 30 precursor ions above a threshold ion count of 3E6 in the MS survey scan with 30.0 s dynamic exclusion. Automatic gain control (AGC) was used to prevent overfilling of the ion trap; 1E5 ions were accumulated for generation of MS/MS spectra. For MS scans, the m/z scan range was 350 to 1500. Five biological replicates of the LC-MS/MS analysis were performed. The raw mass spectral data in our study is available via iProX with identifier PXD022857.

The protein identification and label free quantification were performed through MaxQuant with an integrated Andromeda search engine (v1.5.3.28). Tandem mass spectra were searched against SwissProt Human database (20, 379 sequences) concatenated with reversedecoy database and protein sequences of common contaminants. Trypsin was specified as cleavage enzyme allowing up to 2 missing cleavages, 5 modifications per peptide and 5 charges. Carbamidomethylation on cystine was specified as fixed modification and oxidation on Met, acetylation on protein N-terminal were specified as variable modifications. False discovery rate (FDR) thresholds for protein, peptide and modification site were specified at 0.01. Minimum peptide length was set at 7. All the other parameters in MaxQuant were set to default values. For quantification of the label-free data (intensity ratio calculation), unique + razor peptides were used in protein quantification with 2 minimum ratio count, based on ion intensities of peaks area observed in the LC MS spectra. Proteins that met the expression fold change ≥ 2 for differential levels and *p* value ≤ 0.05 (two-sided unpaired Student’s t-test) with multiple hypothesis testing correction using the Benjamini-Hochberg FDR cutoff of 0.05 were considered for the analyses. The annotation and gene ontology (GO) functional enrichment analysis were performed by Metascape software [29]. Other bioinformatic analysis was performed using Phyper function of the R software package.

### ChIP-seq and bioinformatics

For MTA1 ChIP-seq, crosslinked chromatin complexes were captured from ∼1 × 10^7^ cells same as Metabolic labeling and then sonicated with a Sonics Vcx130pb. The chromatin complexes pre-cleared with vehicle-magnetic beads, and immunoprecipitated with anti-HA-magnetic beads at 4°C for 4 h. The beads were then strictly washed five times at 4°C with low -salt buffer (composition 20 mM Tris-HCl (pH 8.1), 0.1% SDS, 2 mM EDTA, 1% TritonX100, 150 mM NaCl), five times with high-salt buffer prepared as described above but with 500 mM NaCl, and twice with LiCl buffer (10 mM Tris-HCl (pH 8.0), 0.25 M LiCl, 0.5% NP-40, 1% sodium deoxycholate, 1 mM EDTA). The resulting beads were de-crosslinked using Proteinase K. DNA was purified and used for qPCR or next generation sequencing.

Next-generation sequencing libraries were generated and amplified for 15 cycles with BGISEQ kit. 100-300 bp DNA fragments were gel-purified and sequenced with BGISEQ-500 (BGI). Two biological replicates of the ChIP-seq were performed. The raw data are available in the Gene Expression Omnibus database (GSE162932). Raw datasets were mapped to the hg19 human reference genome by Bowtie2 [30]. No more than two mismatches were allowed in the alignment. Reads mapped only once at a given locus were allowed for peak calling. Enriched binding peaks were generated after filtering through control input. Peaks and bigwig files were generated using peak finding algorithm MACS2 software [31]. Sequencing tracks were visualized in Integrative Genomics Viewer (IGV). Genomic distribution, annotation comparison and visualization of O-GlcNAc binding sites was analyzed by ChIPseeker [32].

Heatmaps of sequencing signal density using K-means clustering were generated by deeptools software [33]. The GO functional enrichment analysis was performed by Metascape software [29]. Other bioinformatic analysis was performed using Phyper function of the R software package.

### Quantitative real-time PCR analysis (qPCR)

Total RNA was isolated using the Trizol method (Invitrogen). A total of 5 μg of RNA were reverse transcribed and amplified using One Step SYBR Prime-Script PLUS RT-PCR Kit (TaKaRa) and the Thermal Cycler Dice instrument (TaKaRa) according to the manufacturer’s instructions. The primers of *c-Fos* are sense-5’CTCAAGTCCTTACCTCTTCCG3’; antisense-5’GAGAAAAGAGACACAGACCCAG 3’.

### Luciferase assay

MCF-7/ADR cells were plated at a density of 1 × 10^4^ cells per well in 96-well plates and then co-transfected with *c-Fos*-promoter driven luciferase plasmids and MTA1-WT or MTA1-3A expression plasmids. Cells were treated with DMSO or 100 μM L01 for 48 h, and reporter gene activities were measured with a dual luciferase assay system (Promega, #E1910). The pGL3-basic vector was used as a control.

### RNA sequencing

5 × 10^4^ HA-MTA1-WT or HA-MTA1-3A-transfected MCF-7 cells were collected, washed thrice with ice-cold PBS. RNA extraction was performed using Trizol Reagent (Invitrogen). The rRNA and genomic DNA were removed by MicrobExpress Kit (Ambion) and DNase I (Invitrogen). The RNA was sheared and reverse transcribed with N6 random primers to obtain cDNA library and then qualified by Agilent 2100 Bioanalyzer. Library sequencing was then performed using a combinatorial probe-anchor synthesis (cPAS)-based BGISEQ-500 sequencer (BGI). Two biological replicates of the RNA-seq were performed. The raw data of RNA sequencing is available in the Gene Expression Omnibus database (GSE162932). Raw reads were filtered to remove the adaptor reads. SOAP nuke software was used to remove reads with unknown bases more than 10% and low-quality reads (the ratio of the bases with quality value Q ≤ 15 greater than 50%). Bowtie2 [30] and HISAT software [34] were used to align clean reads to the reference genome hg19. RSEM [35] was then used to quantify the gene-expression level through the fragments per kilobase of transcript per million mapped reads (FPKM) method. The differentially expressed genes (DEGs) were determined by DESeq2 (fold change ≥ 1.5 and false discovery rate (FDR) ≤ 0.05). The GO functional enrichment analysis was performed by Metascape software [29]. Other bioinformatic analysis was performed using Phyper function of the R software package.

### Statistics analysis

Statistical analyses were performed with two-sided unpaired Student’s t-test for single comparison using the GraphPad Prism 8.0 software and Microsoft excel 2019. *p* values < 0.05 was taken as statistically significant. The data are expressed as the means ± SEM. RNA-seq and ChIP-seq were repeated two times (biological replicates). Five biological replicates of the LC-MS/MS analysis were performed.

## Author Contributions

J.Z., YB.L., and X.X. conceived and designed the study. X.X., Q.W., N.Z., and L.W. performed all labeling cell culture experiments and generated samples for LC-MS/MS and sequencing. K.Z., and Q.C. performed LC-MS/MS. YB.L., X.X., YM.L., and W.L. performed all bioinformatics analyze. YB.L., J.Z. and X.X. analyzed data. X.X., YB.L. and J.Z. drafted the manuscript, while all authors reviewed and approved the final version of the manuscript.

## Acknowledgements

This study is supported by the National Natural Science Foundation of China (31971214, 31870793), the National Science and Technology Major Project of China (2018ZX10302205), the Natural Science Foundation of Liaoning Province (2019-MS-042) and the Fundamental Research Funds for the Central Universities (DUT20YG130, DUT20YG116).

## Data availability

All data supporting the findings of this study are available within the article and its supplementary information files. All ChIP-seq and RNA-seq data files along with their associated metadata have been deposited in the GEO database under the accession code GSE162932. The raw mass spectral data in our study is available via iProX with the identifier PXD022857.

