## Supplementary Figure 1;Supplementary Figure 2 for "O-GlcNAc regulates MTA1 transcriptional activity during breast cancer cells genotoxic adaptation"

**Supplementary information**

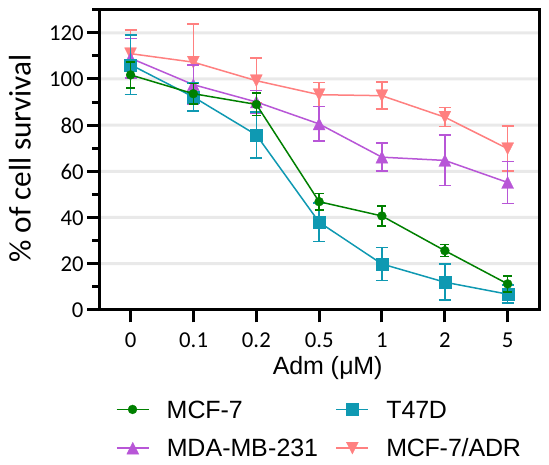

**Supplementary Figure 1** MCF-7, T47D, MDA-MB-231 and MCF-7/ADR cells were treated with the indicated dose of Adm for 48 h. The cell viability was assessed through the CCK8 assay.

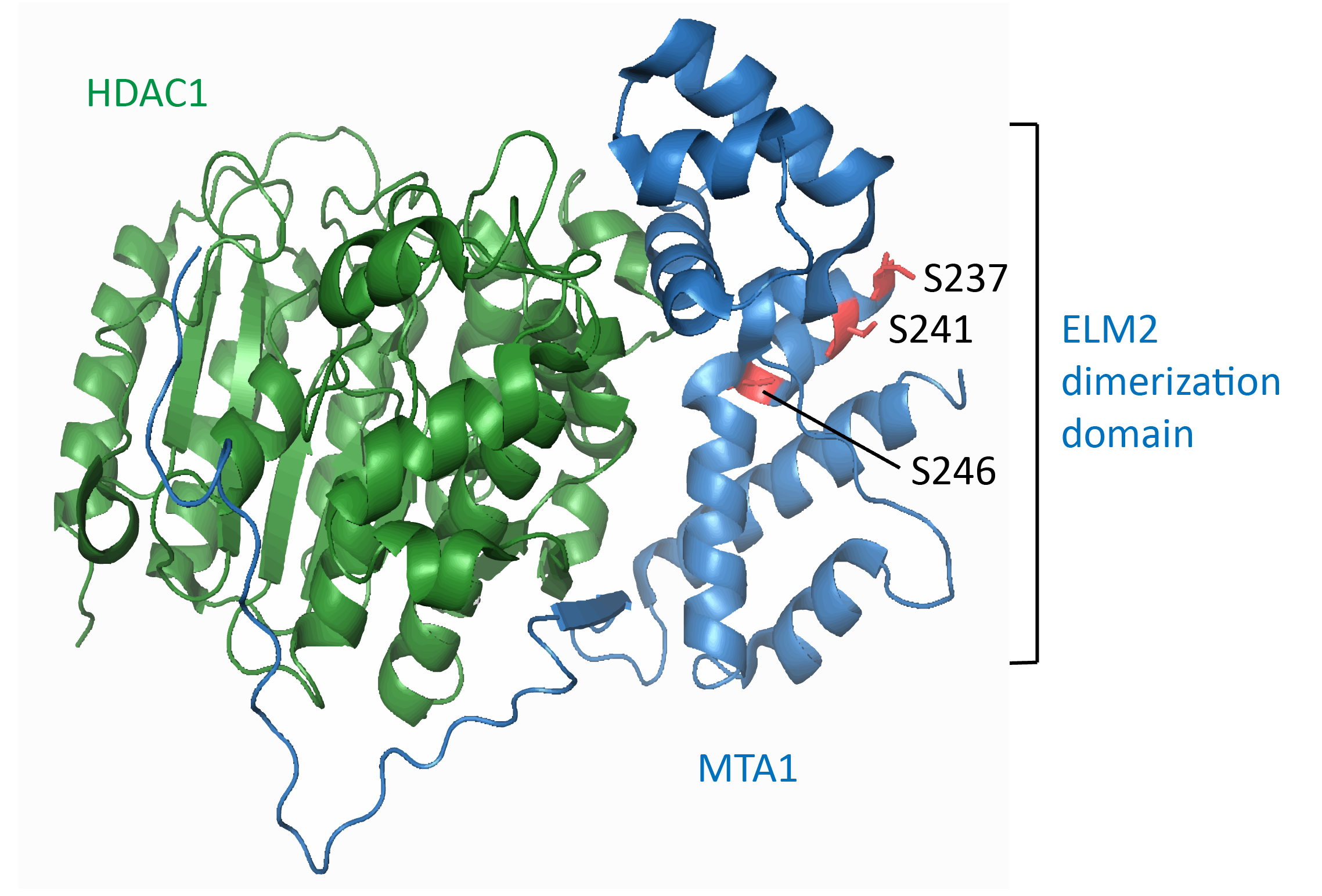

**Supplementary Figure 2** Structure of MTA1 and HDAC1 complex (5ICN). The O-GlcNAc sites of MTA1 are labeled.
